# Alternative splicing use ratio is not conserved between human and mouse immune cells

**DOI:** 10.64898/2026.10.12.758440

**Authors:** Hadas Ner-Gaon, Naama Nudelman, Tal Shay

## Abstract

Most human genes have mouse orthologs. However, orthology at the level of exons and alternative splicing events is not well defined, and the use of alternative splicing events in immune cell types has not been compared between human and mouse.

We mapped exons and alternative splicing events between human and mouse. Based on this mapping, the use ratio of orthologous alternative splicing events was compared between human and mouse immune cell types and fibroblasts.

Out of the 1,823 human-mouse orthologous alternative splicing events that are expressed in the five tested immune cell types, most (62-67%) showed a significant difference in use ratio between human and mouse, and less than third (21-25%) retained the same ratio.

Our results reveal both conserved and divergent alternative splicing usage ratio, and highlight the dynamic evolution of alternative splicing regulation across human and mouse immune cell types. By integrating exon orthology with observed splicing patterns, our analysis provides a functional framework for studying mammalian splicing ratio evolution beyond static transcript annotations.

## 1 Introduction

The mouse is the most commonly used model organism for human immunology (1). However, many studies have repeatedly advised caution when translating findings from mouse to human (1–7), since there are obvious differences between species -life span, size, total number of cell divisions (8), diet, environment, etc. Specifically, there are differences between human and mouse immune systems (9).

In addition to the biological differences between human and mouse immune systems, there are also experimental and ethical considerations that impinge on studies of human and mouse immune systems (3). Studies of human immunology typically focus on cells isolated from peripheral blood, bone marrow, or cord blood, with *in-vivo* studies largely being limited to monitoring clinical manipulations, such as vaccinations or transplantations. In contrast, there are no such limitations on *in-vivo* studies in mice (10). Mouse studies are conducted primarily with inbred mouse strains that are controlled for age, environmental influences, and other confounding factors, in contrast to natural human populations for which these factors cannot always be controlled. Importantly, recent studies have shown that some differences between inbred mice and humans are due to environmental conditions, specifically microbial exposure, and not to species differences. Accordingly, some of the presumed human–mouse differences in immune systems are reduced or disappear when comparing humans to more “natural” mice, such as feral or pet-store mice (11), inbred laboratory mice co-housed with pet-store mice (11), inbred laboratory mice born to wild-type mothers (12), or inbred laboratory mice exposed to common pathogens (13). Still, the many differences between humans and inbred laboratory mice remain a significant impediment to the translation of research from inbred laboratory mice to the clinic.

The known – and yet to be discovered – differences between human and mouse immune systems thus remain one of the major factors contributing to the low success rates of clinical trials. It is estimated that as little as 10-14% of drugs that reach clinical trials receive FDA approval (14, 15). Most of the failures are due to inadequate efficacy or to safety concerns (16), which may derive, at least in part, from differences between model animals and humans. Thus, a better understanding of human–mouse differences is crucial to improving the translation success rate of potential therapeutics.

One of the approaches used to compare the human and mouse immune systems is transcriptional profiling (17), first by microarray and then by RNA sequencing (RNA-seq). We and others have previously shown that microarray-based transcriptional profiles of orthologous immune lineages are globally similar between humans and mice, signatures of lineage specific gene expression are shared, and the expression patterns of most genes are conserved between the two species (18, 19), as well as the transcriptional programs (18). Still, the level of similarity of transcriptional profiles of the human and mouse immune systems has been subject to debate (20–22). A prerequisite for settling this debate is to thus establish a reliable baseline, namely, to accurately map the molecular differences between mouse and human immune systems.

Analysis of RNA-seq datasets can go beyond the gene level into the different transcripts that are transcribed from each gene. Alternative splicing is a major contributor to transcriptomic and proteomic diversity in mammals and is especially prevalent in the immune system (23), where rapid adaptation and cell-type specialization require extensive post-transcriptional regulation. Alternative splicing can alter protein localization, enzymatic activity, interaction domains, or transcript stability, thereby shaping immune-cell identity and function (24). RNA-seq technology has enabled genome-wide studies of transcript diversity. Such studies are now beginning to uncover the abundance and importance of changes in alternative splicing events during immune differentiation and activation (23–25). Few specific alternative splicing events that change use ratio have indeed been well studied (e.g., Ciita (26), Ptprc/Cd45 (27)), but, genome wide, the distribution and regulation of alternative splicing events across the immune system is far from being understood.

Comparative analyses between human and mouse have demonstrated that although many splicing events are evolutionarily conserved, a substantial proportion, estimated at roughly 25–50% of alternative splicing events or isoforms, is species-specific, suggesting ongoing evolutionary rewiring of transcript regulation (28–31). Those large-scale human–mouse splicing comparisons did not study sorted immune cell types and focused on annotated transcript isoforms or computationally reconstructed transcripts (28). However, transcript-level analyses are limited by incomplete transcript annotations, inconsistent isoform catalogue depth between species, and the difficulty of accurately reconstructing full-length transcripts from short-read RNA-seq data. In addition, many annotated transcript variants may represent low-abundance or non-functional products, complicating the interpretation of transcript orthology and functional conservation. These limitations suggest that exon-centered comparative approaches, combined with RNA-seq data, may provide a more robust framework for studying evolutionary conservation of alternative splicing use ratio.

Since the prevalence of species-specific isoforms between human and mouse remains a controversial subject (29–31), we performed a systematic comparative mapping of alternative splicing events between human and mouse using exon-level orthology combined with RNA-seq data from immune cells and fibroblasts. Rather than focusing on transcript orthology, we identified orthologous alternative splicing events and quantified their use ratio directly from splice junctions coverage. This approach enabled us to evaluate the extent to which alternative splicing use ratio is preserved between human and mouse immune cell types. We identified 1,823 human-mouse orthologous alternative splicing events that are expressed in the five immune cell types that were tested. Out of those, 62-67% showed a significant difference in use ratio between human and mouse, and only 21-25% of the orthologous alternative splicing events retained the same use ratio.

## 2 Materials and methods

### 2.1 Mapping of orthologous exons and splicing junctions between human and mouse

Orthologous gene pairs between human and mouse were obtained from BioMart, Ensembl Genes 115 version (32), based on the human genome assembly GRCh38.p14 and the mouse genome assembly GRCm39. For each mouse gene, its human ortholog, homology type and orthology confidence were retrieved.

For each human-mouse high confidence one-to-one pair of orthologous genes, the sequences of all exons of all transcripts were retrieved from Ensembl (32) using Python’s request library (v2.32.4) (https://github.com/psf/requests). For exons that are part of the canonical transcripts, rank (ordinal number) in the canonical transcript was recorded.

In order to identify orthologous exons, for each orthologous gene pair, a pairwise sequence alignment was performed between all human exons and all mouse exons by the PairwieseAligner function of the Biopython library (33), with a global alignment strategy and affine gap penalties. The scoring scheme rewarded nucleotide matches (+2), penalized mismatches (-1), and used higher penalties for gap opening (-12) than gap extension (-4), consistent with standard affine-gap alignment approaches that discourage fragmented insertions/deletions while allowing biologically plausible longer gaps (34). Each exon of an ortholog in one species is aligned to all exons in the ortholog of the other species once. Since the same exon can harbor different untranslated region (UTR) and coding DNA sequence (CDS) boundaries across isoforms, depending on the position of the start or stop codon in each transcript, and since UTRs are less conserved than CDSs (35), the alignment of each exon is only of its CDS portion when available. If the same exon has different coding sequences in different isoforms, its canonical coding sequence is used. If an exon is not in the canonical transcript, the coding sequence of that exon will be taken from the longest coding isoform that contains it, and if no protein-coding transcripts contain it, than the entire exon sequence.

A human exon and a mouse exon that have at least 50% identity to each other are defined as orthologous exons, as previously suggested (35). The start and end coordinates of orthologous exons are defined as orthologous exon boundaries. An exon can have more than one ortholog, so a single human exon boundary may be orthologous to more than one mouse exon boundary (and vice versa). In order to select the best orthologous boundary, all boundaries of the species with multiple orthologous boundaries were ranked using the ratio-based score described below, and the candidate with the highest score was chosen. If two or more candidates were tied for the highest score, the boundary of the canonical exon (rather than a non-canonical exon) was selected. The same procedure is applied in reverse when a single mouse exon boundary has more than one candidate orthologous human boundaries.

The ratio-based score is calculated as the average percent identity of the orthologous exons, multiplied by the ratio of the lower to the higher percent identity between the human and mouse exons, thereby favoring exons pairs with both high overall similarity and balanced conservation across species. A human splicing junction between the end boundary of human exon a and the start boundary of downstream human exon b is defined as orthologous to a mouse splicing junction between the end boundary of mouse exon a’ and the start boundary of downstream mouse exon b’ if exons a and a’ are orthologous, and exons b and b’ are orthologous, and a<b and a’<b’ (Figure 1A).

**Figure 1.**
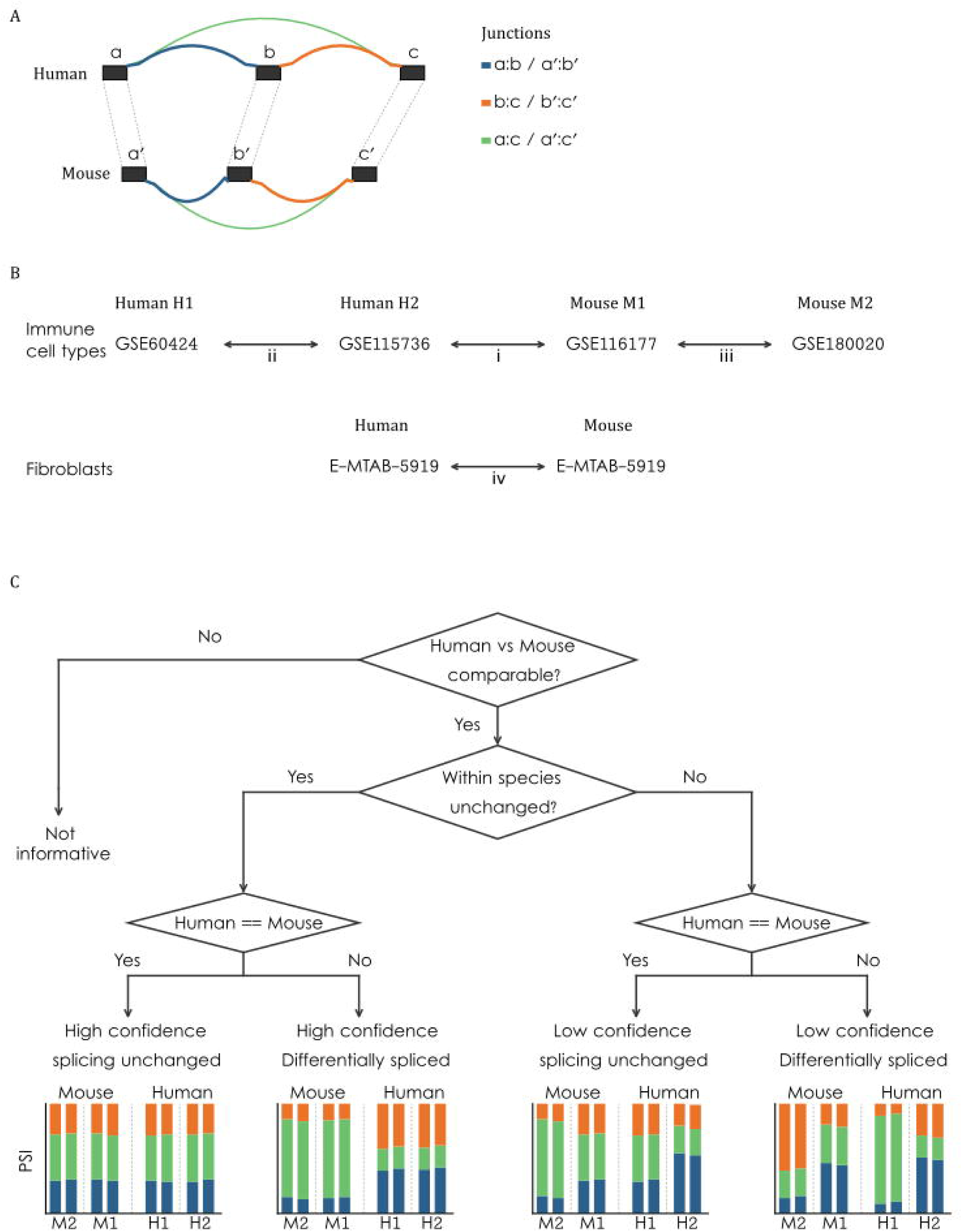
Outline of the human-mouse splicing ratio comparative analysis. A. Definition of orthology for exons, junctions and alternative splicing events for all orthologous genes was performed. Shown is an orthologous skipped exon event in human and mouse. Exons are marked by black boxes and splicing junctions by arcs. Orthologous human and mouse exons are connected by dashed lines, and orthologous junctions are the same color. B. Differential splicing was analyzed for: (i) Each immune cell type between human and mouse datasets, (ii) Each immune cell type between two human studies, and (iii) Each immune cell type between two mouse studies. Additionally differential splicing was analyzed between (iv) human and mouse fibroblasts. C. A flowchart for the decision process of status classification of orthologous alternative splicing events is on the top. Bottom is an illustration representing each event status. Stacked bars show the fraction of junction spanning reads covering each junction participating in the event in each sample. Colors are as in A.

### 2.2 RNA-sequencing datasets

To test for differential splicing between human and mouse, a search was done in GEO (https://www.ncbi.nlm.nih.gov/geo/) (36) to find RNA-sequencing datasets that profiled at least four naive immune cell types: B cells, T cells, natural killer (NK) cells and monocytes (Mono), and include at least seven samples. Specifically, the search query: “b-lymphocytes” [MeSH Terms] AND “t-lymphocytes” [MeSH Terms] AND “killer cells, natural” [MeSH Terms] AND “monocytes” [MeSH Terms] AND (“7”[n_samples]: “100000”[n_samples]) AND “Expression profiling by high throughput sequencing” [Filter] was used. That query on 12/5/2026 returned 17 mouse datasets and 56 human datasets. After filtering out single cell datasets, SuperSerieses, datasets without naïve immune cells, datasets with low depth, and datasets with less than four cell types, two mouse datasets and six human datasets remained. One mouse and one human dataset that were profiled by the same lab, GSE115736 and GSE116177 (37), were chosen for the Human-Mouse comparison, to minimize technical differences. For the Human-Human comparison, GSE115736 samples were compared to GSE60424 samples (38), as this dataset is deep enough and of high quality. For the Mouse-Mouse comparison, GSE116177 samples were compared to the other mouse dataset that remained following the filtering - GSE180020 (39). Human-Mouse fibroblasts RNA-seq was taken from E-MTAB-5919 dataset (40) that includes dermal fibroblasts from both human and mouse. All datasets have at least two replicates in all cell types. For the selected datasets, raw sequencing reads were downloaded from the NCBI Sequence Read Archive (SRA) (41) using fastq-dump from the NCBI SRA Toolkit (v3.0.0, Available from: https://github.com/ncbi/sra-tools). Hisat2 (42) (version 2.0.5) was used to align human or mouse RNA-seq reads to the Homo sapiens genome (GRCh38) or Mus musculus genome (GRCm39), respectively. The resulting binary alignment map (BAM) files were sorted and indexed by SAMtools (43) (version 1.9). To evaluate the quality of the data, the flagstat option of SAMtools was used. In all datasets, all samples had at least 17 million properly mapped reads and more than 90% of the reads were mapped (Tables S1-S6). Thus, all samples were used for the analysis.

### 2.3 Defining human-mouse orthologous alternative splicing events

Splice junctions were extracted from all BAM files using LeafCutter (v0.2.9) (44), which identifies splice junctions from mapped junction spanning RNA-seq reads. The ‘prepare_junctions.py’ script of LeafCutter was modified to output all the Junction Spanning Reads (JSR) and their counts from each dataset, without clustering. Since the RNA-seq datasets used are not stranded, the parameter ignore_strand was used. This parameter makes it possible to merge junction spanning reads (JSR) counts across strands.

To enable differential splicing analysis between human and mouse, only orthologous splicing junctions were used. In order to create the combined human and mouse JSR expression table required for differential splicing analysis by LeafCutter, each mouse junction was assigned with the coordinates of its human orthologous junction in the JSR expression table that was served as input of LeafCutter.

To ensure robust detection and cross-dataset comparability, only orthologous junctions that fulfil the following criteria were included in the analysis: (1) Junction must be covered in each of the four datasets being compared by at least one read, and (2) junction must be covered by at least 10 reads in at least two samples from two different datasets. These stringent criteria minimize spurious junctions and increase the likelihood of sufficient evidence across datasets. The remaining junctions were assigned to LeafCutter clusters. A LeafCutter cluster is a set of junctions that share start or end coordinates, and it represents an alternative splicing event. Junctions that were not assigned to clusters, that is, junctions that do not share start or end genomic locations with other junctions, were removed.

### 2.4 Differential splicing analysis

Using the JSR expression table, differential splicing analysis was performed according to the workflow of LeafCutter (44) (https://davidaknowles.github.io/leafcutter/articles/Usage.html). H. sapiens genome annotation file (GCF_000001405.39_GRCh38.p13_genomic.gff) was downloaded from NCBI (https://www.ncbi.nlm.nih.gov/genome/51) and parsed to fit the LeafCutter input format. Default parameters were used, except for –g and -i. The -g parameter (min_samples_per_group) sets the minimal number of samples in each group that are covered by at least min_coverage reads. The -i parameter (min_samples_per_intron) sets the minimum number of samples in which an intron must have at least one supporting read to be retained; introns that do not meet this threshold are excluded. Both parameters are assigned a number which is 60% of the number of samples in the smallest group in the relevant comparison (2-8, Table S1).

The fraction of JSRs of a junction in each LeafCutter cluster in each sample is termed the percent spliced in (PSI). Differential splicing is measured in terms of the change in PSI between conditions, ΔPSI. Max(|ΔPSI|) was defined as the maximal abs(ΔPSI) value in each comparison, in each LeafCutter cluster.

### 2.5 Classification of Alternative Splicing Events

Based on the differential splicing analysis, in each cell type, each comparable alternative splicing event (success LeafCutter cluster) was assigned a status - Differentially spliced: p.adjust ≤ 0.05 and max(|ΔPSI|) ≥ 0.1); Splicing unchanged: p.adjust > 0.05 or max(|ΔPSI|) < 0.05; Not informative: Event that is not differentially spliced or unchanged. Alternative splicing events that were informative (either differentially spliced or splicing unchanged) in the human-mouse comparison were assigned a high confidence level if both within species comparisons were splicing unchanged, and low confidence otherwise (Figure 1B,C). In fibroblast, confidence level was not assigned as there is a single dataset in each species.

### 2.6 Functional Enrichment Analysis

Functional enrichment analyses were performed using gseapy enrich module (45) with three MSigDB (46) gene set collections: GO_Biological_Process_2025 (c5.go.bp), Reactome_2022 (c2.cp.reactome) and Hallmark (h.all; MSigDB) version v2024.1.Hs, with a fixed background of 1,408 unique gene symbols corresponding to the genes that contain the 1,823 comparable alternative splicing events. Two cell-type-independent lists were defined by pooling status calls across all five immune cell types: DS_only (genes differentially spliced in at least one cell type and never called splicing-unchanged in any cell type; n = 670) and SU_only (genes containing events that are classified as splicing-unchanged in at least one cell type and not classified as differentially spliced in any cell type; n = 173). Enrichment was tested for each of the lists against each of the three gene-set libraries, using the shared 1,408-gene background. Benjamini–Hochberg False Discovery Rate (FDR) correction (47) was applied per annotation source. Enrichment results were considered statistically significant at FDR ≤ 0.1.

## 3 Results

### 3.1 The human-mouse exon and alternative splicing events orthology map

There are 16,167 high confidence human-mouse one-to-one orthologous genes, out of which 15,127 have at least two exons in each species. Considering all exons from all the transcripts of each orthologous gene, those genes in human include 724,579 exons, out of which 691,489 exons (95%) have mouse orthologs. In mouse, those 15,127 genes contain 392,376 exons, out of which 379,935 exons (97%) have human orthologs. The higher number of human exons with mouse orthologs is due to the larger number of exons in human (48, 49). There are 377,955 unique orthologous exons boundaries. Orthologous junctions are defined as junctions that share start and end orthologous exons boundaries.

We have used two human and two mouse RNA-seq datasets which profile multiple immune cell types, B cells, CD4 T cells, CD8 T cells, natural killer cells, and monocytes (37–39), later termed B, T4, T8, NK and MO, respectively, to compare alternative splicing use ratio between human and mouse in the immune system.

There are 110,174 orthologous junctions that are covered by RNA-seq reads in all four RNA-seq datasets. From those junctions, 87,461 (79%) are covered by at least 10 reads in at least two samples in at least two datasets. Most of those junctions do not represent alternative splicing events. However, 4,962 junctions (6%) form 1,823 alternative splicing events for which more than one transcript is expressed in human and mouse immune cells (Table S7). Those 1,823 alternative splicing events are mapped to 1,408 unique genes and include 967 skipped exons, 392 alternative 3’ splice site, 355 alternative 5’ splice site, and 109 complex events that do not fall into one of the above categories (Table S8).

The alternative splicing events were tested in each immune cell type for differential splicing between the human datasets GSE60424 (H1) and GSE115736 (H2), between the mouse datasets GSE116177 (M1) and GSE180020 (M2), and between human dataset H2 and mouse dataset M1 (Figure 1B), and classified in each comparison as not-comparable, not informative, splicing unchanged or differentially spliced (Figures 1C, 2).

### 3.2 Most alternative splicing events change their ratio between human and mouse immune cells

In the Human-Human comparisons of all immune cell types, the splicing of most comparable (that is, covered by enough reads to estimate the splicing ratio) alternative splicing events is unchanged (54-61%, Table 1, Figures 2A, S1A). Similarly, in the Mouse-Mouse comparisons, most events are classified as unchanged or not informative (Figures 2B, S1B). In contrast, in the Human-Mouse immune cells comparison, more than half of the comparable alternative splicing events in each cell type are differentially spliced (62-67%, Figures 2C, S1C).

**Figure 2.**
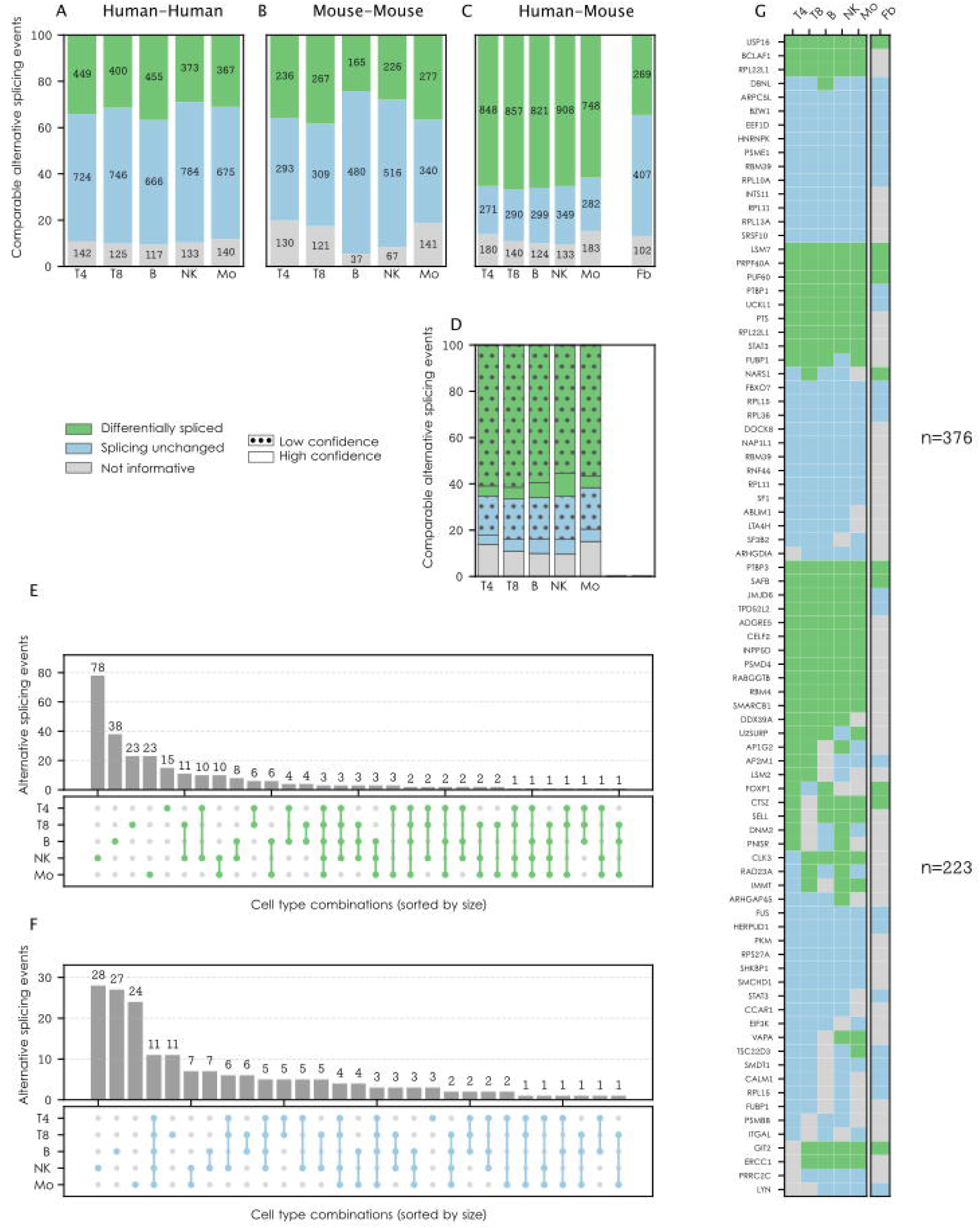
Classification of human-mouse orthologous alternative splicing events based on the conservation of their use ratio. Stacked bars display the percentages of orthologous alternative splicing events which are differentially spliced, use ratio unchanged, or not informative in the (A) human-human comparison, (B) mouse-mouse comparison and (C) human-mouse comparison. D. Same as B, with low confidence events marked by dots. E. UpSet plot of high confidence shared differentially spliced events between cell types. F. UpSet plot of high confidence shared events whose splicing ratio was unchanged. G. Heatmap of high confidence differntially spliced or unchanged splicing events across immune cell types and fibroblasts. Only events which are differntially spliced or unchanged with high confidence in at least three immune cell type are displayed. Events are sorted by number of cell types in which they have high confidence. Gene symbols are shown for the containing genes. Fb – fibroblasts.

**Table 1:**
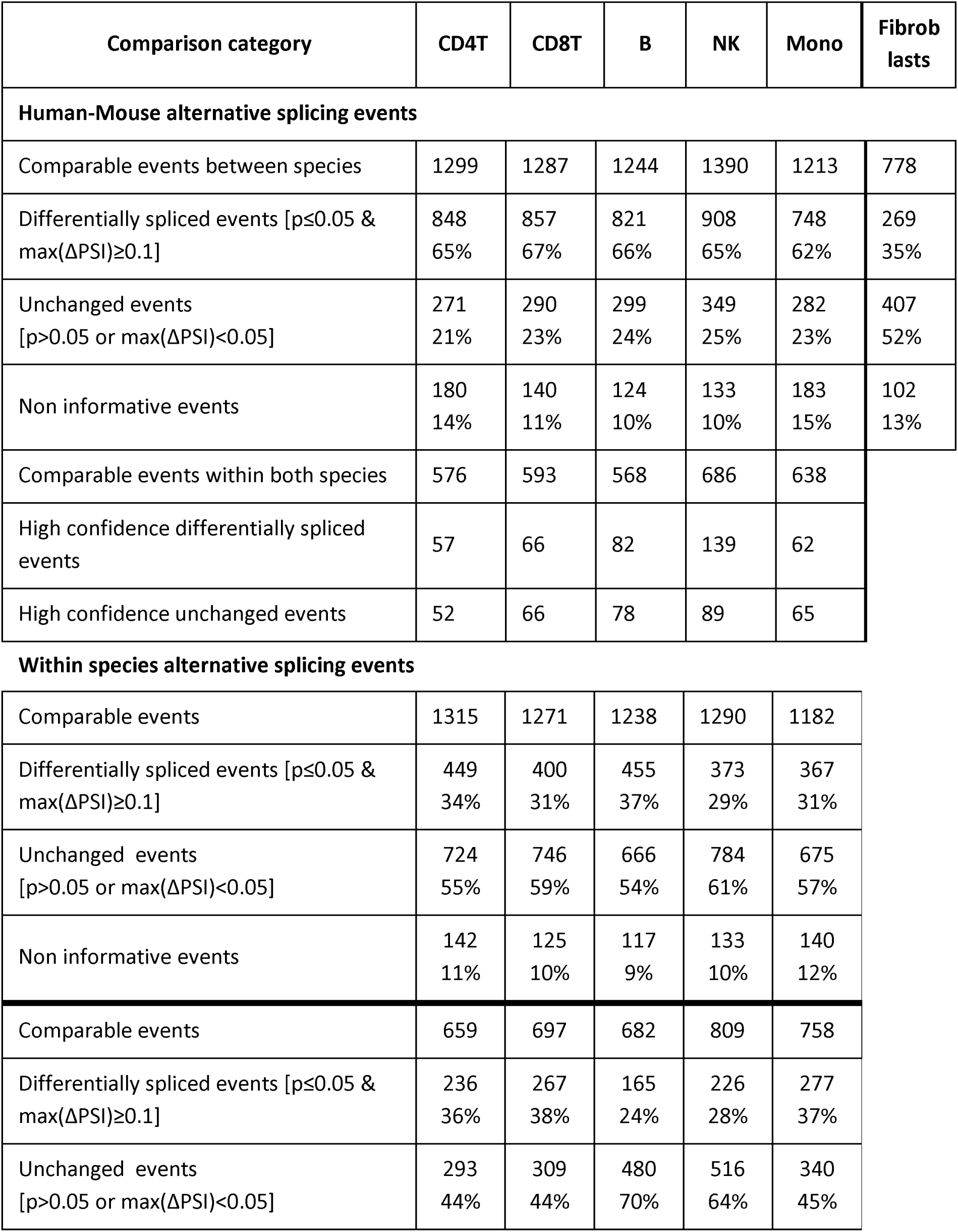
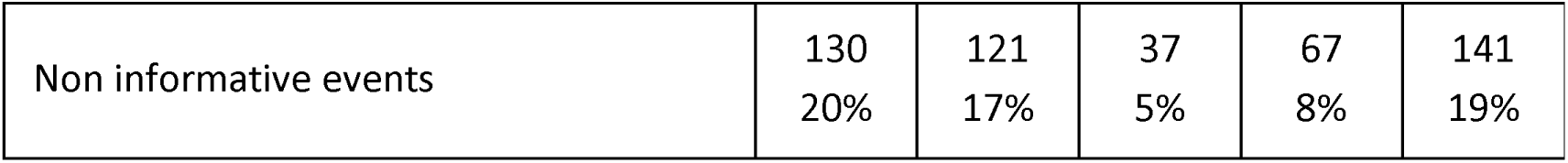
Output of the differential splicing analysis.

| Comparison category | CD4T | CD8T | B | NK | Mono | Fibroblasts |
| --- | --- | --- | --- | --- | --- | --- |
| **Human-Mouse alternative splicing events** |  |  |  |  |  |  |
| Comparable events between species | 1299 | 1287 | 1244 | 1390 | 1213 | 778 |
| Differentially spliced events [$p \leq 0.05$ & $\max(\Delta\text{PSI}) \geq 0.1$] | 848<br>65% | 857<br>67% | 821<br>66% | 908<br>65% | 748<br>62% | 269<br>35% |
| Unchanged events [$p > 0.05$ or $\max(\Delta\text{PSI}) < 0.05$] | 271<br>21% | 290<br>23% | 299<br>24% | 349<br>25% | 282<br>23% | 407<br>52% |
| Non informative events | 180<br>14% | 140<br>11% | 124<br>10% | 133<br>10% | 183<br>15% | 102<br>13% |
| Comparable events within both species | 576 | 593 | 568 | 686 | 638 |  |
| High confidence differentially spliced events | 57 | 66 | 82 | 139 | 62 |  |
| High confidence unchanged events | 52 | 66 | 78 | 89 | 65 |  |
| **Within species alternative splicing events** |  |  |  |  |  |  |
| Comparable events | 1315 | 1271 | 1238 | 1290 | 1182 |  |
| Differentially spliced events [$p \leq 0.05$ & $\max(\Delta\text{PSI}) \geq 0.1$] | 449<br>34% | 400<br>31% | 455<br>37% | 373<br>29% | 367<br>31% | |
| Unchanged events [$p > 0.05$ or $\max(\Delta\text{PSI}) < 0.05$] | 724<br>55% | 746<br>59% | 666<br>54% | 784<br>61% | 675<br>57% | |
| Non informative events | 142<br>11% | 125<br>10% | 117<br>9% | 133<br>10% | 140<br>12% |  |
| Comparable events | 659 | 697 | 682 | 809 | 758 |  |
| Differentially spliced events [$p \leq 0.05$ & $\max(\Delta\text{PSI}) \geq 0.1$] | 236<br>36% | 267<br>38% | 165<br>24% | 226<br>28% | 277<br>37% | |
| Unchanged events [$p > 0.05$ or $\max(\Delta\text{PSI}) < 0.05$] | 293<br>44% | 309<br>44% | 480<br>70% | 516<br>64% | 340<br>45% | |

|  |  |  |  |  |  |
| --- | --- | --- | --- | --- | --- |
| Non informative events | 130<br>20% | 121<br>17% | 37<br>5% | 67<br>8% | 141<br>19% |

The relatively high fraction of within species differential splicing in human (29-37%) and mouse (24-38%) immune cells (Table 1) indicates that the identification of differential splicing is either noisy or affected by the sequencing protocol, and possibly the different composition of subtypes also contributes to the difference. Thus, we further classified the differentially spliced and unchanged events between species into high and low confidence, depending on the within species comparison of each event. As the human and mouse datasets that were not directly compared (H1 and M2) had lower average read number per sample compared to H2 and M1 (Table S1), the requirement for within species comparison prevented all events which were not comparable in one of the within species comparison from reaching high confidence, and thus the number of alternative splicing events that can potentially be of high confidence is roughly 50% of the number of alternative splicing events that can be compared between species (Table 1, rows ‘Comparable events between species and ‘Comparable events within both species’). Thus, out of the 748-908 human-mouse differentially spliced events, only 7-15% are of high confidence, and out of the 271-349 human-mouse alternative splicing events whose splicing is unchanged, 19-26% are of high confidence (Figures 2D, S1D, Table S9). We consider this to be an underestimation, as in addition to events which cannot be of high confidence due to low coverage, many events are more different between species than within species, as is the case for PUF60 in monocytes (Figure S2),

### 3.3 Low confidence human-mouse differential splicing events are shared between immune cell types

Most of the 270 differential splicing events that are classified as high confidence are only of high confidence in one immune cell type (177 events, 66%, Figures 2E, S1E, Table S10), with NK cells leading with the largest number of unique events (78), followed by 38 events in B cells. Only three events are shared between all five immune cell types – USP16 (Figure 3), RPL22L1 (Figure S3), and BCLAF1 (Figure S4). In contrast, when considering also the low confidence differential splicing events, 1027/1,278 (80%) are differentially spliced in more than one immune cell type, and 415 (32%) are differentially spliced in all immune cell types tested (Figure S1E, compare to Figure 2E, Table S10). The genes containing differential splicing events are not enriched for any functional term.

**Figure 3.**
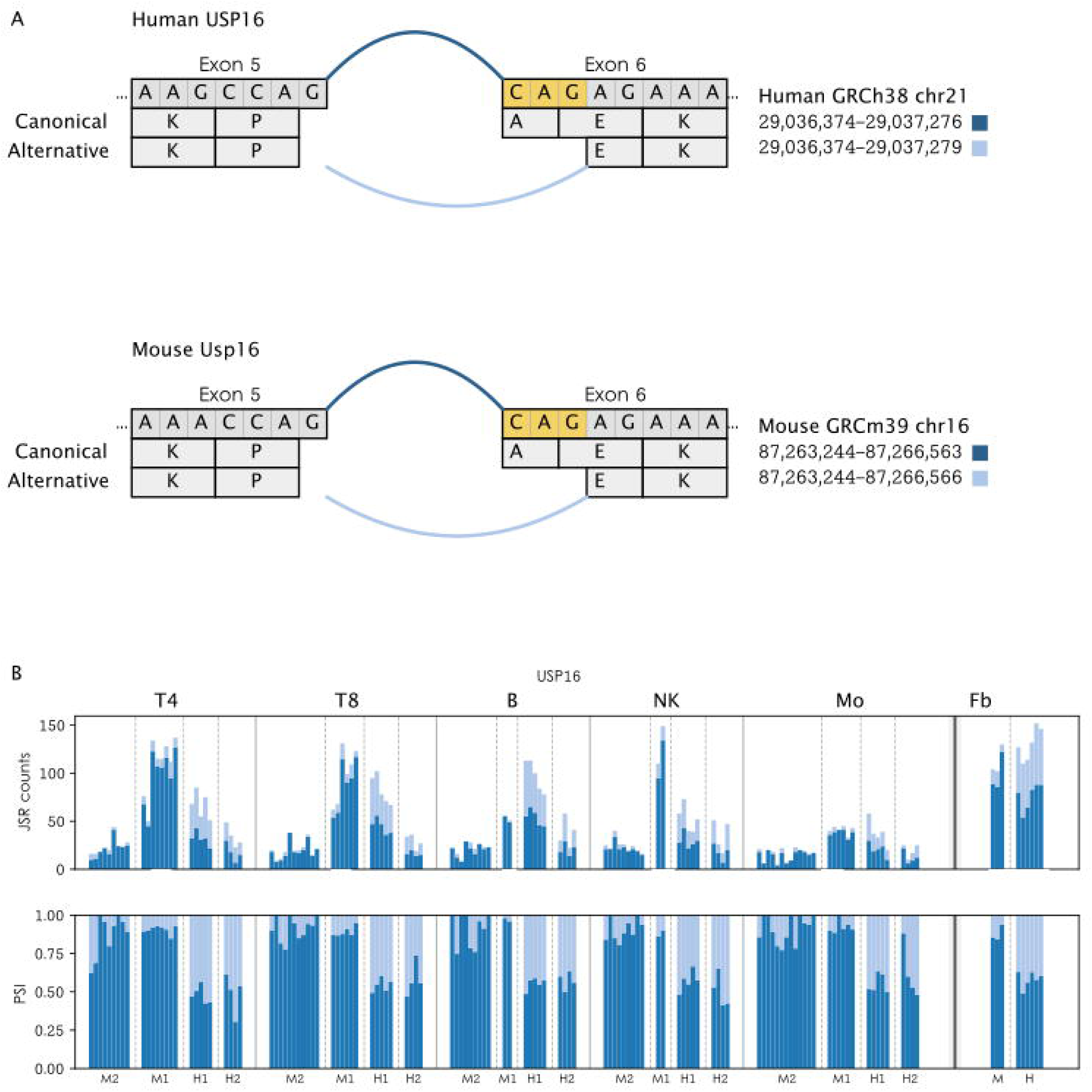
The orthologous alternative 3’ splicing event in USP16 gene is differentially spliced between Human and Mouse. A. An orthologous alternative 3’ splicing event of exon 6. The transcript that includes the blue junction codes for an additional Alanine at position 150, in the human canonical transcript, compared to the transcript that includes the light blue junction, which is shorter by three nucleotides. B. Stacked bars display the number of junction spanning reads that cover the alternative splicing junctions in each sample. Colors are as in A. C. Same as B, numbers are normalized to the total to display PSI values.

### 3.4 The human-mouse alternative splicing events that do not change ratio are shared between immune cell types

There are 181 alternative splicing events that display the same ratio in human and mouse in at least one cell type with high confidence (Table S10). Out of those, 28, 27 and 24 events are unchanged with high confidence in only NK cells, B cells and monocytes, respectively. There are 11 alternative splicing events that do not change their splicing ratio across the five immune cell types with high confidence - PSME1, RPL13A, INTS11, RPL11, SRSF10, RBM39, BZW1, RPL10A, EEF1D, ARPC5L, and HNRNPK (Figure 2F, Figure 4). Similarly to differential splicing events, when considering also low confidence splicing unchanged events, 63 events are shared between all immune cell types tested. However, the most prevalent splicing unchanged group is events that are unchanged only in NK cells (Figure S1F).

**Figure 4.**
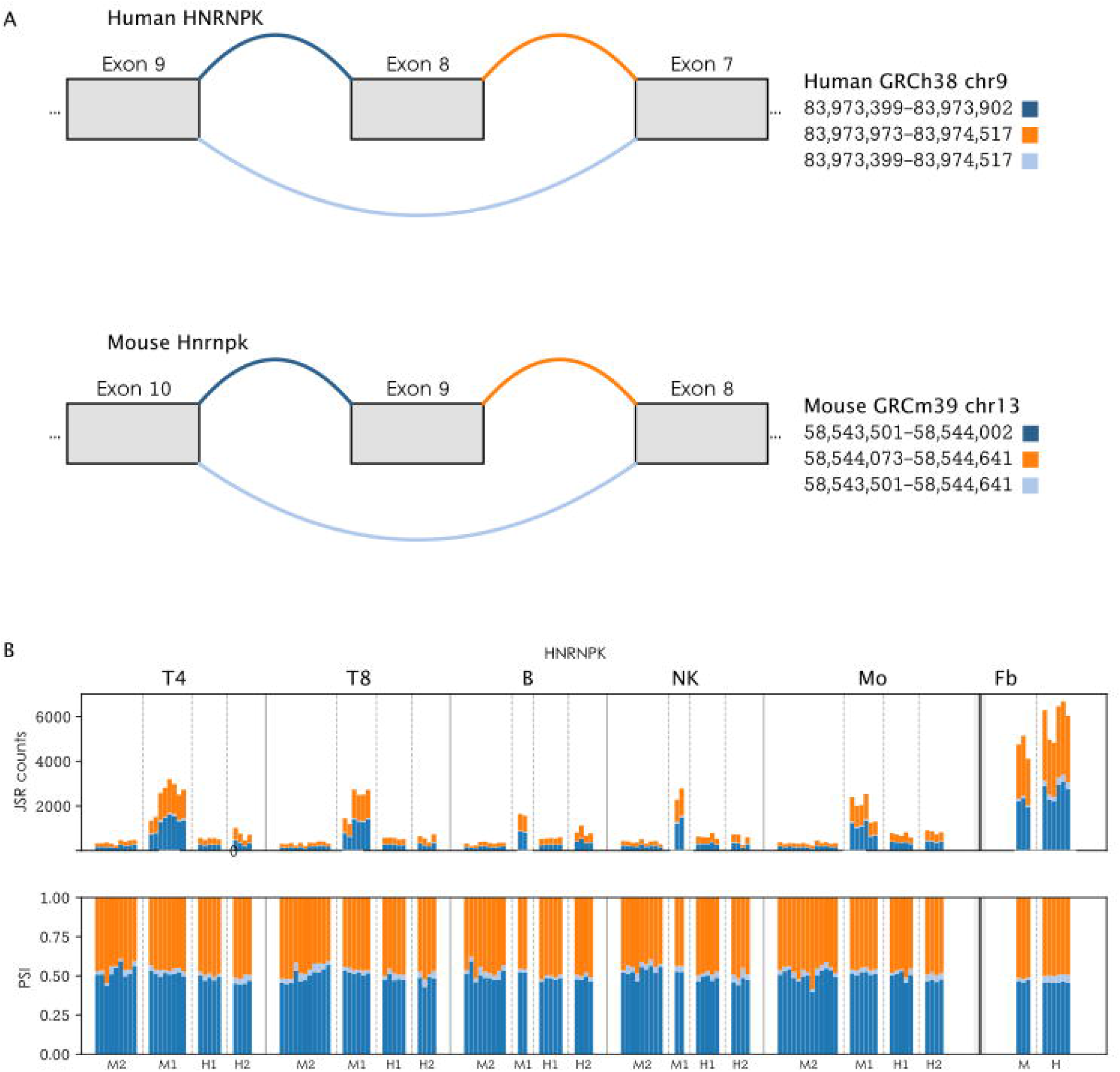
The orthologous skipped exon event in HNRNPK gene displays the same splicing ratio in human and mouse. A. An orthologous skipped exon event. Skip of 72bp, a.a 111-134 (exon 8 in human and exon 9 in mouse). B. Stacked bars display the number of junction spanning reads that cover the alternative splicing junctions in each sample. Colors are as in A. C. Same as B, numbers are normalized to the total to display PSI values.

The list of splicing unchanged genes in any of the cell types (n = 173), despite being shorter than that of the differentially spliced genes, was significantly enriched for 11 Reactome terms (FDR ≤ 0.1) related to translation- and ribosome-associated biology (Table S11). A set of nine ribosomal proteins (RPL3, RPL11, RPL13A, RPL10A, RPL37A, RPL36, RPL15, FAU, RPS27A) was annotated with all 11 significant terms, including seemingly unrelated categories such as “Nervous System Development” and “Signaling by ROBO Receptors”.

### 3.5 Most differential splicing events are unique to the immune system

Human-mouse differential splicing events can be due to a difference in splicing code between human and mouse, in which case such events will be found in all cell types that express the gene, or due to a change in splicing regulation, in which case they may be found only in some cell types. As we found many human-mouse differential splicing events that are shared between immune cell types, at least with low confidence, we wanted to extend beyond the immune system to test which of these explanations is more likely for each human-mouse differential splicing event. Thus, we compared the splicing ratio between immune cells and an RNA-seq dataset that profiled human and mouse dermal fibroblasts (40). Out of the 1,823 human-mouse orthologous alternative splicing events that were comparable in immune cell types, 57% were not comparable due to not being expressed at all or high enough in fibroblasts in one or both species (Figures S6, S7). Out of the 778 comparable events (43%) between human and mouse fibroblasts, 269 (35%) were differentially spliced, 407 (52%) were unchanged, and 102 (13%) were not informative (Figure 2B). Out of the 351–415 differentially spliced events in immune cell types, 141–163 (37–41%) are differentially spliced also in fibroblasts, 161–197 (46–51%) are unchanged, and 43–58 (11–14%) are not informative (Figure 5A). Out of the 117–158 events whose splicing is unchanged in immune cell types, 85–110 (65–75%) are unchanged also in fibroblasts, 14–30 (11–21%) are differentially spliced, and 13–21 (10–14%) are not informative (Figure 5B). Conversely, out of the 269 differentially spliced events in fibroblasts, 141–163 (81–85%) are also differentially spliced in immune cell types, only 14–30 (8–17%) are unchanged in immune cell types, and 4–11 (2–7%) are not informative (Figure 5C). Out of 407 events whose splicing is unchanged in fibroblasts, 161–197 (52–60%) are differentially spliced in immune cells, 85–110 (25–32%) do not change splicing in immune cell types, and 36–54 (11–18%) are not informative (Figure 5D).

**Figure 5.**
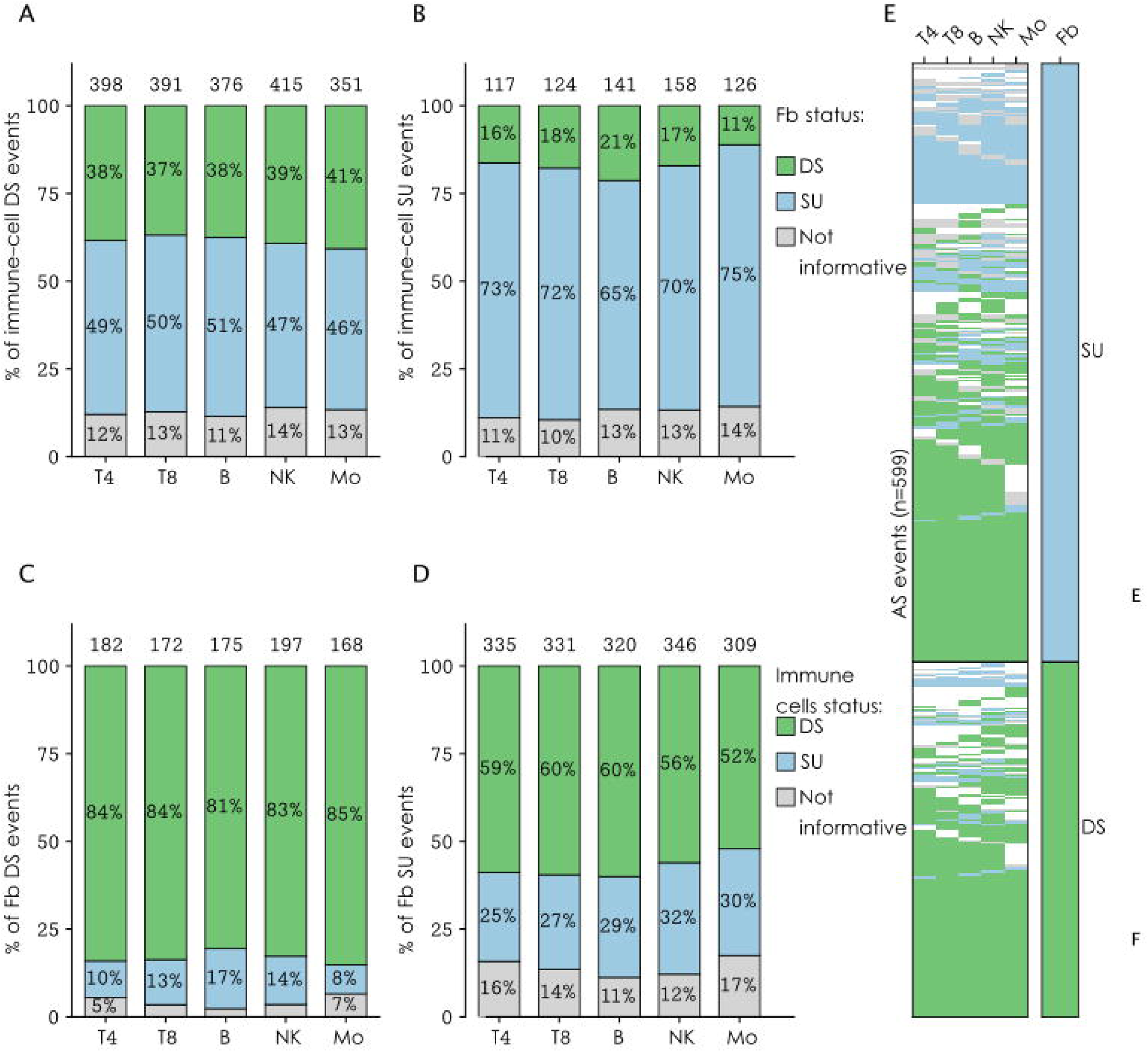
Comparison of human-mouse splicing differences in the immune system and in fibroblasts. Distribution of the status in fibroblasts of alternative splicing events that are (A) differentially spliced or (B) their splicing status is unchanged between human and mouse, in each immune cell type. Distribution of the status in immune cells of alternative splicing events that are (C) differentially spliced or (D) their splicing is unchanged between human and mouse fibroblasts, shown separately for each immune cell type. Number above each bar represents that number of alternative splicing events that are comparable in the immune cell type and in fibroblasts. (E) Heatmap of the splicing status in each immune cell type (CD4T, CD8T, NveB, NK, Mono) and in fibroblasts, for all alternative splicing events with a DS/SU status in fibroblasts (n = 599). Each row represents one alternative splicing event, ordered by the number of immune cell types in which it is differentially spliced. DS – differentially spliced; SU – splicing unchanged.

## 4 Discussion

Here, we performed a systematic, exon-level comparison of alternative splicing between human and mouse across five naïve immune cell types and dermal fibroblasts, using orthologous exon, junction and alternative splicing events mapping combined with RNA-seq splice-junction quantification. We identified 1,823 expressed orthologous alternative splicing events, the majority of which (62-67% per cell type) were differentially spliced between species, while a minority displayed similar use ratio in human and mouse. These findings indicate that orthology of gene sequence, expression level pattern or alternative splicing repertoire does not guarantee orthology at the level of alternative splicing usage, with implications for how mouse models are interpreted in immunological research. In contrast to immune cells, fibroblasts display largely conserved splicing patterns between species for the same alternative splicing events, suggesting that the observed changes are tissue specific, and not due to a change in splicing code.

Most human-mouse orthologous alternative splicing events displayed different splicing ratio in the cell types in which they were comparable. This is in contrast to the within-species comparisons, in which most of those events displayed conserved splicing ratio. While this difference indicates that the human-mouse prevalent difference in splicing ratio is not merely due to technical differences between RNA-seq datasets, it also means that there are many events that change due to technical differences. Thus, we limit the interpretation of differentially spliced events only to high confidence events, that is, events that displayed conserved splicing ratio within species, reducing the number of alternative splicing events displaying differential splicing from 1,278 to 270.

Only three alternative splicing events displayed high confidence ratio change between human and mouse in all five immune cell types, and all were alternative 3’ splicing events in which two AG dinucleotides are one or four nucleotides away, which affect three or six nucleotides, respectively. In USP16 (Figure 3), a histone H2A deubiquitinase, the three additional nucleotides code for an additional Alanine at the beginning of exon 6. Cysteine 205 that is required for USP16 deubiquitinase activity (50) is located towards the end of exon 6, and might be affected by the change. In the ribosome-associated protein RPL22L1 (Figure S3), the three additional nucleotides add Glutamine. In the transcriptional repressor BCLAF1 (Figure S4), the six additional nucleotides turn Arg-Ser-Arg (R-S-R) into Ser-Ser-Arg-Ser-Arg (S-S-R-S-R). Genes containing NAGNAG events, that is, AG dinucleotides that are separated by one nucleotide, are conserved between human and mouse and were found to be enriched for DNA binding genes (51). In all three events described above, the shorter exon was more frequent in human immune cells. However, testing all human-mouse comparable NAGAG events, the fraction of differential splicing was not significantly different from that of all comparable events, and there was no preferential use of the shorter exon by either species (data not shown).

As stated above, naturally most within-species comparisons displayed conserved splicing ratio, but only the minority of human-mouse events displayed a conserved splicing ratio – 611 events in 531 genes, and if only considering high confidence, only 181 events of conserved splicing in 162 genes. The genes carrying the conserved ratio alternative splicing events were enriched for translation related mechanisms, mostly due to ribosomal proteins. An example of a gene that exhibits a high confidence conserved splicing ratio across species in all cell types tested is exon 8 skipping of the heterogeneous nuclear ribonucleoprotein K, HNRNPK (Figure 4), which codes for the linker between the second and third KH nucleic acid binding domain. As both upregulation and downregulation of HNRNPK have been identified in cancer (52), it is possible that an accurate ratio between its isoforms is strongly selected for. Indeed, of the four orthologous alternative splicing events in HNRNPK, three are classified as unchanged splicing ratio, and one is low confidence differential splicing in the last exon. Genes encoding core RNA-processing and splicing machinery are generally among the most evolutionarily constrained genes in mammals, in terms of sequence and expression level. This is consistent with stronger purifying selection and intolerance to loss of function compared to genes involved in more peripheral or context-dependent signaling pathways, such as those in immune and inflammatory response, which show greater divergence in both sequence and expression across mammals (53). Thus, it is possible that those genes are also under stronger purifying selection for splice ratio. If this is the case, core genes are expected to display splicing ratio conservation in all tissues and cell types.

To test if the human-mouse conservation level of splicing ratio is similar for non-immune cells, we compared immune cell types to a purified but non-immune cell type - fibroblasts. In all immune cell types, most comparable alternative splicing events were differentially spliced between human and mouse. In contrast, the same events mostly displayed unchanged splicing in fibroblasts (Figure 5A). An example is the skipped exon in UCKL1, which is skipped often in mouse immune cells, but is hardly skipped in human immune cells or fibroblasts of both species (Figure S8). However, some events are differentially spliced in immune cells and fibroblasts, for example USP16 (Figure 3) and PUF60 (Figure S2).

Out of the splicing unchanged events of each immune cell type, only ∼20% were differentially spliced in fibroblasts, consistent with core genes being unchanged in all cell types (Figure 5B). Similarly, ∼80% of the differentially spliced events in fibroblasts were also differentially spliced in each immune cell type (Figure 5C), but only 50-60% of the splicing unchanged events (Figure 5D). Thus, there is a set of core genes that contain comparable alternative splicing events whose splicing ratio is similar in human and mouse immune cells and fibroblasts and is enriched for core cellular processes – HNRNPK, RPL10A, RPL15, RPL3, RPL36, RPL37A, SRSF5, SRSF7 and more.

Several alternative splicing events whose ratio is up-regulated in cancer were also detected in normal immune cells, with a tendency toward higher ratio in mouse naïve immune cells compared to their human counterparts: exon 9 skipping in ZNF207, which promotes hepatocellular carcinoma and is regulated by hnRNPA1 (54), exon 5 skipping in PUF60 (Figure S2), which is increased in multiple cancers (55) and is regulated by SAP155/SF3b1 (56). This suggests that so-called “tumor-associated” splice isoforms are not cancer-specific but rather reflect broader regulatory programs that can be differentially utilized across physiological and species contexts. As cancer poses a much stronger selective force for humans than for short-lived mouse, it is tempting to assume that cancer promoting isoforms will tend to be higher in mouse. However, there are also examples in which the oncogenic splicing variants display higher ratio in humans, as is the case for ATG16L1 (57) and FLNA (58).

Though we have not explicitly studied differential splicing between immune cells, we saw previously reported events, including the skipped exon of DOCK8 (Figure S5) (59) and the differentially spliced skipped exons of PTPRC (60) (Figure S6). In both human and mouse, CD47 has five different tails due to the skipping combinations of the later exons, but their ratio differ between species (Figure S7) (61).

Whereas most human exons have mouse orthologs, and most human genes have more than one transcript, only 1823 alternative splicing events were covered by enough RNA-seq reads to be compared between the species. This means that in order to accurately compare splicing ratio between species, mapping of orthologous alternative splicing events between species should be done per cell type.

The ability to identify comparable orthologous alternative splicing events is limited to relatively highly expressed genes, as it requires a minimal coverage of the splicing junctions by RNA-seq reads. Thus, only 1823 alternative splicing events were covered by enough RNA-seq reads to be compared between the species in at least one cell type, and those events are within only 1,408 genes. While 1096 genes have a single event, 314 genes carry 2-6 orthologous alternative splicing events, which often display both splicing difference and splicing unchanged. HNRNPK discussed above carry five orthologous alternative splicing events – two skipped exons with unchanged splicing ratio, two alternative 5’ splice site (one unchanged and one not comparable) and one alternative 3’ splice site which is differentially spliced. Much deeper sequencing is required to compare splicing ratio of most orthologous alternative splicing events.

Our findings emphasize that understanding which aspects of mouse biology are comparable to human biology is critical when using mouse models to study human physiology and disease. While orthologous genes are often assumed to perform similar functions across species, our results demonstrate that orthology should also be considered at the transcript and alternative splicing levels rather than solely at the gene level. In many cases, the relative usage of orthologous alternative splicing events was not conserved between human and mouse, despite conservation of the corresponding genes. These differences in splicing ratio may have important functional consequences, as distinct isoforms of the same gene can differ in protein structure, localization, stability, interaction partners, or regulatory properties.

Several technical and biological factors may have influenced both the number and identity of the detected differential and conserved alternative splicing events between human and mouse orthologous genes. First, sequencing depth directly affects the number of testable alternative splicing events between and within species. Only 1823 events were comparable in at least one cell type, and even fewer events were comparable in both human-mouse and within species comparisons, limiting the potential number of high confidence findings. Second, the number of biological replicates affect statistical power, and in some datasets and cell type there were only two replicates. Third, sequencing-related biases may differ between datasets due to differences in library preparation protocols, read quality, read length, and alignment efficiency. Fourth, biological heterogeneity also represents an important source of variation. Human samples inherently contain substantial genetic variation between donors, which may increase inter-individual variability in splicing patterns compared to the isogenic mouse strain used here. Furthermore, some annotated cell populations likely contain distinct cellular subpopulations with unique transcriptional and splicing programs, potentially masking or diluting species-shared splicing signatures. Additional factors such as differences in sampled tissues (Spleen in one mouse dataset, multiple tissues in the other mouse dataset, and peripheral blood in human), developmental stage/age (5-12 weeks in mouse vs adults in human), and sex composition between datasets (not reported in both mouse datasets and GSE115736, only females in GSE60424) may further contribute to species-specific splicing patterns and complicate the interpretation of conserved versus divergent use ratio of alternative splicing events. Fifth, RNA-sequencing displays sequence dependent coverage bias (62). This bias is canceled out for within species differential splicing, but between species comparison of splicing junctions spanned by different sequences with different bias may inflate the estimate of differentially spliced events.

To partially account for some of those sources of variability and improve the robustness of our analysis, we classified differentially spliced and splicing conserved events into two confidence levels. High-confidence events were defined as events in which the splicing pattern was consistently conserved within each species across independent comparisons (human-to-human and mouse-to-mouse), supporting the reproducibility of the observed regulation. In contrast, low-confidence events included cases in which such within-species conservation cannot be estimated due to low coverage, or was not observed, suggesting that these events may be more sensitive to technical variability, sample heterogeneity, or context-specific regulation.

The prevalence of human-mouse differential splicing identified in this study may contribute to the limited translational relevance observed for some mouse models of human conditions. Together, these results highlight the importance of incorporating alternative splicing-level analyses into comparative and translational studies to better evaluate the extent to which mouse models accurately recapitulate human molecular biology.

## 5 Statements

### Data availability statement

Publicly available datasets were analyzed in this study. This data can be found here: https://www.ncbi.nlm.nih.gov/geo/query/acc.cgi?acc=GSE115736, https://www.ncbi.nlm.nih.gov/geo/query/acc.cgi?acc=GSE116177, https://www.ncbi.nlm.nih.gov/geo/query/acc.cgi?acc=GSE60424, https://www.ncbi.nlm.nih.gov/geo/query/acc.cgi?acc=GSE180020.

Programs can be found in: https://github.com/Tal-Shay-Group/Human_Mouse

### Author contributions

HN-G performed the analysis. NN performed the initial analysis. TS supervised the research. HN-G and TS wrote the manuscript. All authors approved the manuscript.

## Acknowledgments

We sincerely thank all members of the ImmGen consortium - it has been a genuine pleasure and privilege, scientifically and personally, to be part of ImmGen. We would also like to express our gratitude to Mr. Yehuda Barouch for his continuous support.

