## Supplementary material for "Alternative splicing use ratio is not conserved between human and mouse immune cells": Sup_Material: Table 1.DOCX

### 1 Supplementary Tables

Table S1: Datasets summary

|  |  |  |  | Human | | Human | | Mouse | | Mouse | |
| --- | --- | --- | --- | --- | --- | --- | --- | --- | --- | --- | --- |
|  | -i, -g | | | GSE115736 | | GSE60424 | | GSE116177 | | GSE180020 | |
|  | H_M | H_H | M_M | Average mapped | #Replicates | Average mapped | #Replicates | Average mapped | #Replicates | Average mapped | #Replicates |
| CD4T | 3 | 2 | 4 | 19,078,819 | 5 | 41,438,696 | 4 | 41,722,693 | 8 | 29,128,326 | 9 |
| CD8T | 3 | 2 | 3 | 18,848,484 | 5 | 35,718,782 | 4 | 41,214,305 | 6 | 28,183,087 | 11 |
| B | 2 | 2 | 2 | 18,524,465 | 5 | 44,352,197 | 4 | 37,062,398 | 2 | 28,573,763 | 9 |
| NK | 2 | 2 | 2 | 19,224,036 | 5 | 31,677,199 | 4 | 45,056,630 | 2 | 29,482,658 | 9 |
| Mono | 3 | 2 | 3 | 18,940,852 | 5 | 34,966,894 | 4 | 35,132,845 | 6 | 27,270,447 | 13 |
|  |  |  |  | Human |  | Mouse |  |  |  |  |  |
|  |  |  |  | E-MTAB-5919 | | | |  |  |  |  |
| Fibroblasts | 2 |  |  | 52,002,728 | 6 | 48,274,978 | 3 |  |  |  |  |

#-i,"--min_samples_per_intron", default=5, Ignore introns used (i.e. at least one supporting read) in fewer than n samples

#-g,"--min_samples_per_group", default=3, Require this many samples in each group to have at least min_coverage reads

Table S2: Immune cells that were used from Human GSE115736 dataset

| Cell name | Cell type | Tissue | Replicates | Reads | Reads length |
| --- | --- | --- | --- | --- | --- |
| CD4T | T | Peripheral Blood | 5 | single | 100bp |
| CD8 | T | Peripheral Blood | 5 | single | 100bp |
| Mono | Monocyte | Peripheral Blood | 5 | single | 100bp |
| NK | NK | Peripheral Blood | 5 | single | 100bp |
| NveB | Naïve B | Peripheral Blood | 5 | single | 100bp |

Table S3: Immune cells that were used from Mouse GSE116177 dataset

| Cell name | Cell type | Tissue | Replicates | Reads | Reads length |
| --- | --- | --- | --- | --- | --- |
| CD4 | Total CD4+ T | Spleen and Lymph nodes | 2 | single | 100bp |
| EffCD4T | CD4 Effector T | Spleen and Lymph nodes | 2 | paired | 200bp |
| MemCD4T | CD4 Memory T | Spleen and Lymph nodes | 2 | paired | 200bp |
| NveCD4T | CD4 Naive T | Spleen and Lymph nodes | 2 | paired | 200bp |
| CD8T | Total CD8+ T | Spleen and Lymph nodes | 2 | single | 100bp |
| MemCD8T | CD8+ Memory T | Spleen and Lymph nodes | 2 | paired | 200bp |
| NveCd8T | CD8+ Naive T | Spleen and Lymph nodes | 2 | paired | 200bp |
| BCell (PtB) | Total B Cells | Peritoneal Cavity Lavage | 2 | single | 100bp |
| MonoPB | Total Monocyte | Peripheral Blood | 2 | single | 100bp |
| MonoBM | Total Monocyte | Bone Marrow | 2 | single | 100bp |
| InfMono | Inflammatory Monocyte | Bone Marrow | 2 | single | 100bp |
| NK | Mature NK | Spleen | 2 | paired | 200bp |

Table S4: Immune cells that were used from Human GSE60424 dataset

| Cell name | Cell type | Tissue | Replicates | Reads | Reads length |
| --- | --- | --- | --- | --- | --- |
| B | B-cells | Peripheral Blood | 4 | Paired | 100bp |
| CD4 | T-cells | Peripheral Blood | 4 | Paired | 100bp |
| CD8 | T-cells | Peripheral Blood | 4 | Paired | 100bp |
| MO | Monocytes | Peripheral Blood | 4 | Paired | 100bp |
| NK | NK | Peripheral Blood | 4 | paired | 100bp |

Table S5: Immune cells that were used from Mouse GSE180020 dataset

| Cell name | Cell type | Tissue | Replicates | Reads | Reads length |
| --- | --- | --- | --- | --- | --- |
| B | B Follicular | Spleen | 9 | Paired | 76bp |
| CD4 | Naïve T | Spleen | 9 | Paired | 76bp |
| CD8 | Effector memory T | Spleen | 11 | Paired | 76bp |
| Monocyte | Monocyte | Spleen | 13 | Paired | 76bp |
| NK | Natural Killer | Spleen | 9 | Paired | 76bp |

Table S6: Compared samples

| Cell type | Human GSE115736 | Human GSE60424 | Mouse GSE116177 | Mouse GSE180020 |
| --- | --- | --- | --- | --- |
| CD4 | CD4T | CD4 | CD4T, EffCD4T,  MemCD4T, NveCD4T | Naïve T |
| CD8 | CD8T | CD8 | CD8T, MemCD8T,  NveCd8T | Effector memory T |
| B | NveB | B | BCell | B Follicular |
| NK | NK | NK | Mature NK | Natural Killer |
| Monocyte | Mono | MO | MonoPB, MonoBM,  InfMono | Monocyte |

Table S9: High-confidence differentially spliced (DS) and splicing-unchanged (SU) events

| Cell type | DS total | DS high confidence | DS high confidence % | SU total | SU high confidence | SU high confidence % |
| --- | --- | --- | --- | --- | --- | --- |
| CD4T | 848 | 57 | 6.7% | 271 | 52 | 19.2% |
| CD8T | 857 | 66 | 7.7% | 290 | 66 | 22.8% |
| NveB | 821 | 82 | 10.0% | 299 | 78 | 26.1% |
| NK | 908 | 139 | 15.3% | 349 | 89 | 25.5% |
| Mono | 748 | 62 | 8.3% | 282 | 65 | 23.0% |

Table S11**.** Reactome_2022 pathway enrichment in the splicing-unchanged (SU) gene list

| Term | Fold Enrichment | Gene Names^1^ | Number of Genes | FDR |
| --- | --- | --- | --- | --- |
| Eukaryotic Translation Elongation | 7.5 | **RPL37A, RPL36, FAU, RPL13A, RPL15, RPL11, RPL10A, RPL3**, EEF2**, RPS27A** | 10 | 0.022 |
| SRP Dependent Cotranslational Protein Targeting To Membrane | 6.84 | **RPL37A, RPL36**, TRAM1**, FAU, RPL13A, RPL15, RPL11, RPL10A, RPL3, RPS27A** | 10 | 0.028 |
| Response Of EIF2AK4 GCN2 To Amino Acid Deficiency | 6.74 | **RPL37A, RPL36, FAU, RPL13A, RPL15, RPL11, RPL10A, RPL3, RPS27A** | 9 | 0.042 |
| Cellular Response to Starvation | 6.15 | **RPL37A, RPL36, FAU, RPL13A, RPL15, RPL11, RPL10A, RPL3, RPS27A** | 9 | 0.058 |
| Influenza Infection | 5.82 | **RPL37A, RPL36, FAU, RPL13A, RPL15, RPL11, RPL10A, RPL3**, XPO1**, RPS27A** | 10 | 0.042 |
| rRNA Processing | 4.75 | **RPL37A, RPL36, FAU, RPL13A, RPL15, RPL11, RPL10A, RPL3**, RIOK3**, RPS27A** | 10 | 0.089 |
| Eukaryotic Translation Initiation | 5.06 | **RPL37A, RPL36, FAU, RPL13A, RPL15, RPL11, RPL10A, RPL3**, EIF3K**, RPS27A** | 10 | 0.075 |
| Cellular Responses To Stimuli | 2.54 | SOD2**, RPL36**, COX7A2L**, RPL13A, RPL15**, HERPUD1**, RPL37A**, HDGF**, RPL11, RPL3**, XPO1**, FAU**, MAPK14, ELOC, COX6B1**, RPL10A, RPS27A**, PRDX5, PSMB8, PSME1, CAPZB, NCF4, KAT5 | 23 | 0.089 |
| Nervous System Development | 3.12 | PFN1**, RPL36**, CLASP1**, RPL13A, RPL15**, CLASP2**, RPL37A, RPL11, RPL3**, PRKCQ**, FAU**, ELOC**, RPL10A**, PLCG1, SDCBP**, RPS27A**, PSMB8, PSME1, ZSWIM8, PTPRA, PIK3CD | 21 | 0.036 |
| Signaling By Robo Receptors | 5.65 | PFN1**, RPL37A**, PSMB8, PSME1**, RPL36**, CLASP1, ZSWIM8**, FAU**, ELOC**, RPL13A, RPL15**, CLASP2**, RPL11, RPL10A, RPL3, RPS27A** | 16 | 0.005 |
| Regulation Of Expression Of Slits And Robos | 5.54 | **RPL37A**, PSMB8, PSME1**, RPL36**, ZSWIM8**, FAU**, ELOC**, RPL13A, RPL15, RPL11, RPL10A, RPL3, RPS27A** | 13 | 0.022 |

^1^In bold, genes shared between all enriched terms

### 2 Supplementary Figures Legends

**Figure S1 - Classification of human-mouse orthologous alternative splicing events based on the conservation of their use ratio.** Stacked bars display the percentages of orthologous alternative splicing events which are differentially spliced, use ratio unchanged, not informative or not comparable in the (A) human-human comparison, (B) mouse-mouse comparison and (C) human-mouse comparison. Percentages are out of orthologous alternative splicing events. D. Same as C, with low confidence events marked by dots. E. UpSet plot of low and high confidence shared differentially spliced events between cell types. F. UpSet plot of low and high confidence events whose splicing ratio was unchanged. G. Heatmap of high confidence shared differentially spliced or unchanged splicing events across immune cell types and fibroblasts. Only events which are differentially spliced or unchanged with high confidence in two immune cell types are displayed. Events significant in more than two immune cell types are displayed in Figure 2G. Gene symbols are shown for the containing genes.

**Figure S2** **- The orthologous skipped exon 5 event in PUF60 gene displays differential splicing between human and mouse**. A. The schematics of the orthologous skipped exon event. Exons are represented by rectangles, splicing junctions by arcs. B. Stacked bars display the number of junction spanning reads that cover the alternative splicing junctions in each sample. Colors are as in A. C. Same as B, numbers are normalized to the total to display PSI values. Fb – fibroblasts.

**Figure S3** - **The orthologous alternative 3' splicing event in RPL22L1 exon 2 is differentially spliced between human and mouse.** A. An orthologous alternative 3' splicing event of exon 2. The transcript that includes the blue junction codes for an additional Glutamine, in the human canonical transcript, compared to the transcript that includes the light blue junction, which is shorter by three nucleotides. B. Stacked bars display the number of junction spanning reads that cover the alternative splicing junctions in each sample. Colors are as in A. C. Same as B, numbers are normalized to the total to display PSI values.

**Figure S4 -** **The orthologous alternative 3' splicing event in BCLAF1 exon 4 is differentially spliced between human and mouse.** A. An orthologous alternative 3' splicing event of exon 4. The transcript that includes the blue junction codes for an additional Serine and Arginine, in the human canonical transcript, compared to the transcript that includes the light blue junction, which is shorter by six nucleotides. B. Stacked bars display the number of junction spanning reads that cover the alternative splicing junctions in each sample. Colors are as in A. C. Same as B, numbers are normalized to the total to display PSI values.

**Figure S5 - The orthologous alternative first exon in DOCK8 gene displays the same splicing ratio in human and mouse but differential splicing between immune cells.**

A. An orthologous alternative first exon. The transcript that includes the blue junction codes for longer transcript, in compare to the transcript with the light blue junction. The mouse transcript with the light blue junction biotype is protein coding, CDS not defined. B. Stacked bars display the number of junction spanning read that support the alternative junctions in each sample. Colors are as in A. C. Same as B, numbers are normalized to the total to display PSI values.

**Figure S6 - The orthologous complex skipped exons event in PTPRC displays differential splicing between immune cell types and species.** A. The schematics of the orthologous skipped exons event. Exons are represented by rectangles, splicing junctions by arcs. B. Stacked bars display the number of junction spanning read that support the alternative junctions in each sample. Colors are as in A. C. Same as B, numbers are normalized to the total to display PSI values.

**Figure S7 - The orthologous complex skipped exons event in CD47 displays differential splicing between species, but is similar between immune cell types.** A. The schematics of the orthologous skipped exons event. Exons are represented by rectangles, splicing junctions by arcs. B. Stacked bars display the number of junction spanning read that support the alternative junctions in each sample. Colors are as in A. C. Same as B, numbers are normalized to the total to display PSI values.

**Figure S8 - The orthologous skipped exon event in UCKL1 displays differential splicing between human and mouse, but is unchanged in fibroblasts.** A. The schematics of the orthologous skipped exon event. Exons are represented by rectangles, splicing junctions by arcs. B. Stacked bars display the number of junction spanning read that support the alternative junctions in each sample. Colors are as in A. C. Same as B, numbers are normalized to the total to display PSI values.
