## Supplementary figures and images for "Alternative splicing use ratio is not conserved between human and mouse immune cells"

### Image 1.TIFF

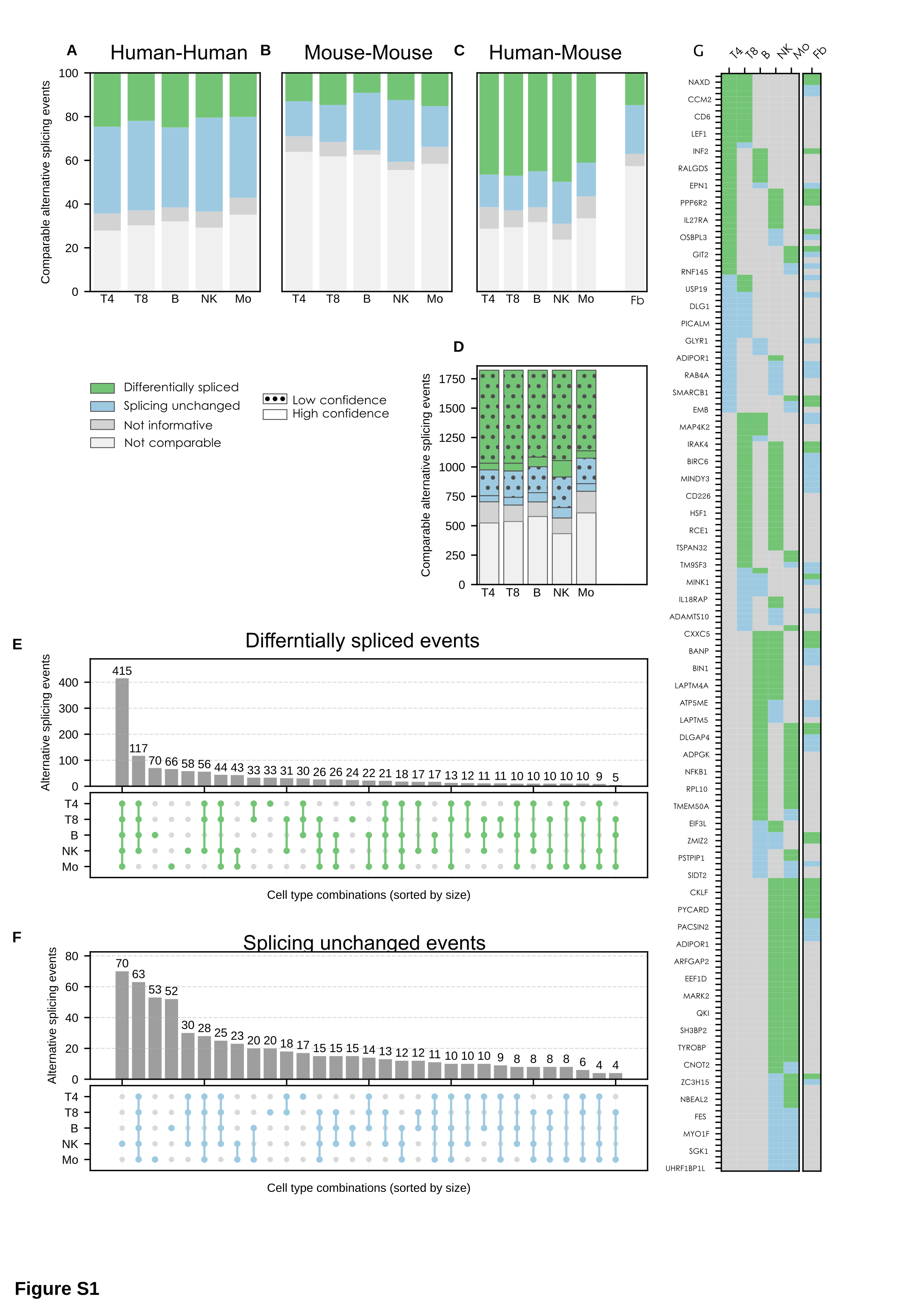

### Image 2.TIFF

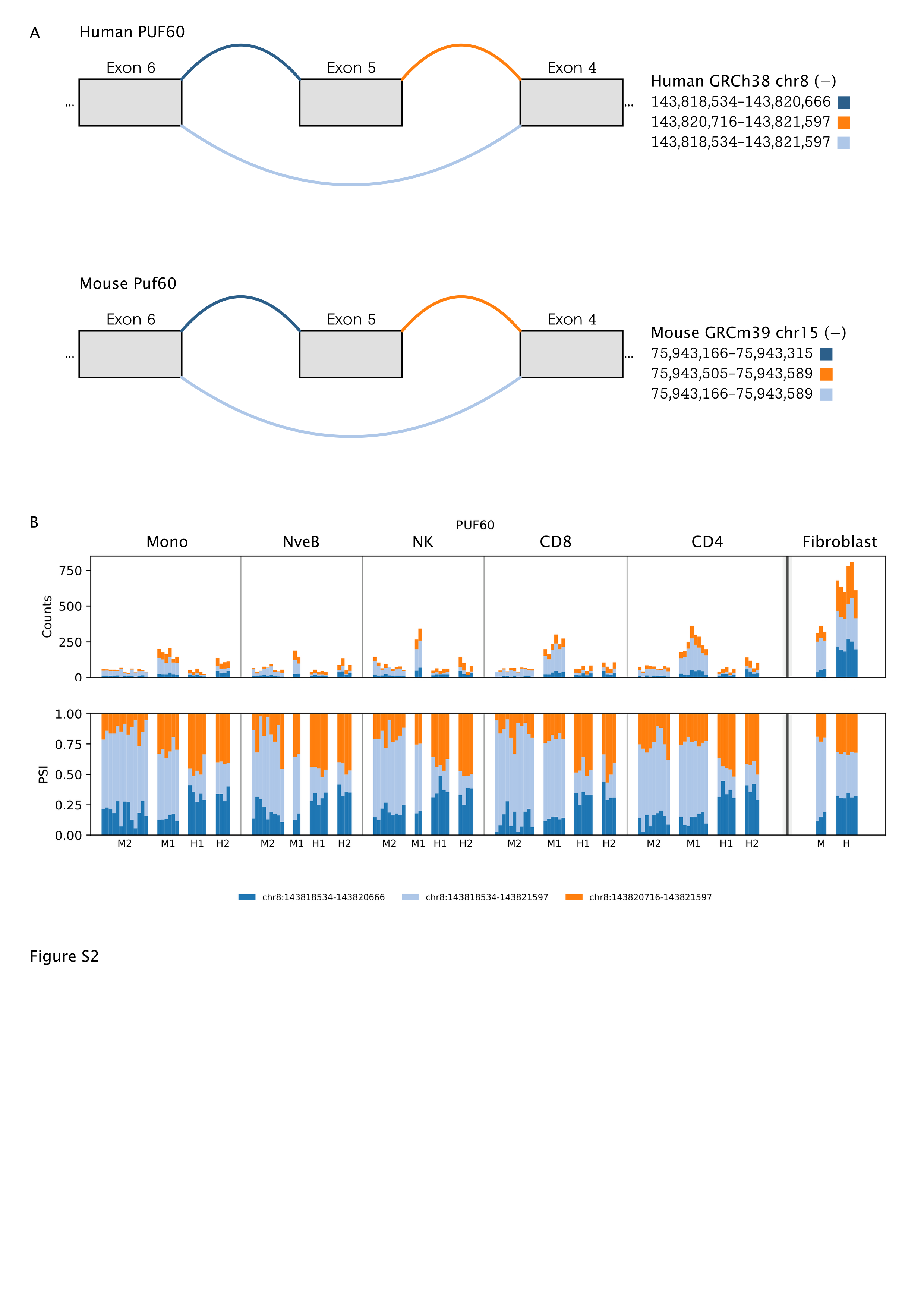

### Image 3.TIFF

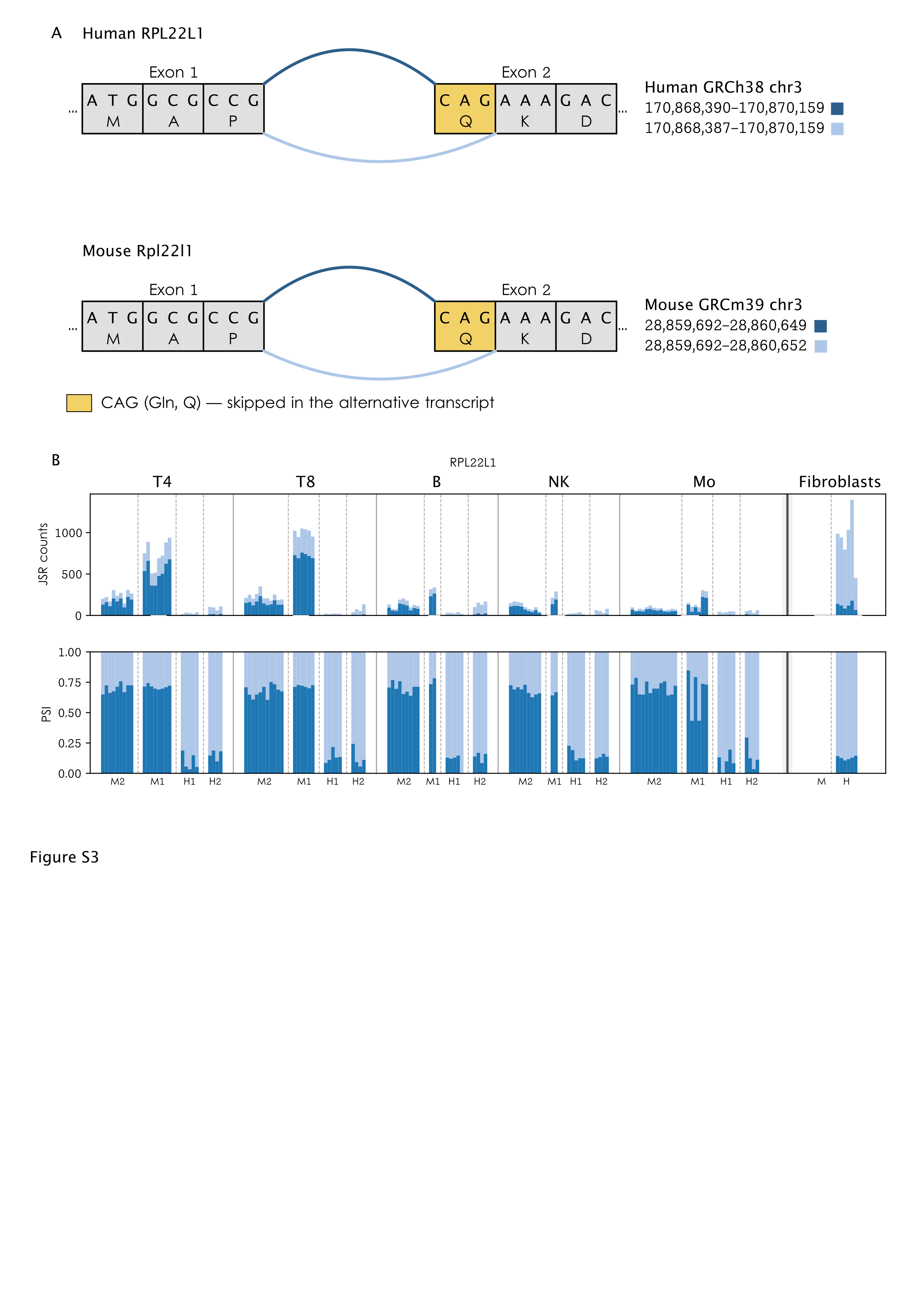

### Image 4.TIFF

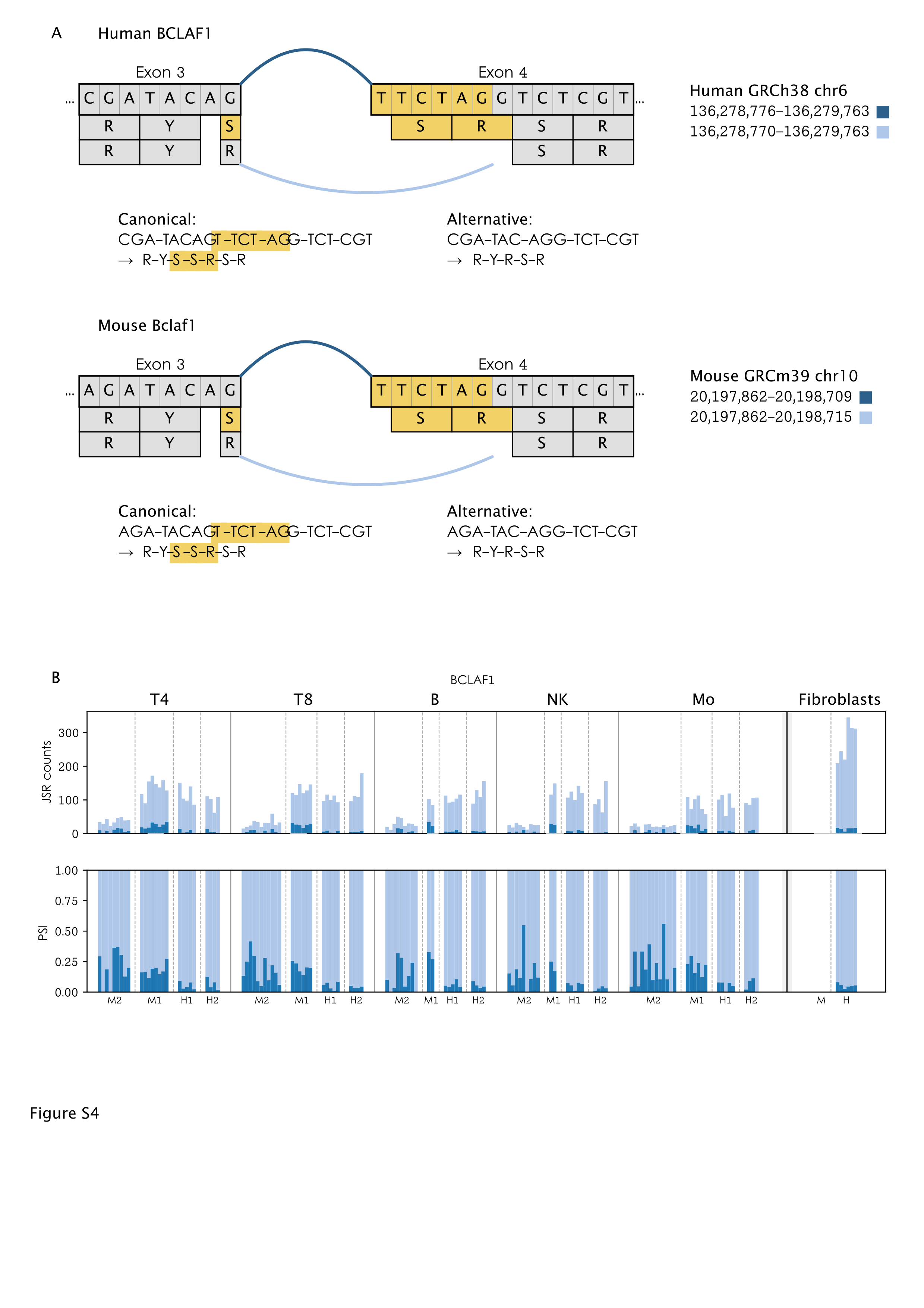

### Image 5.TIFF

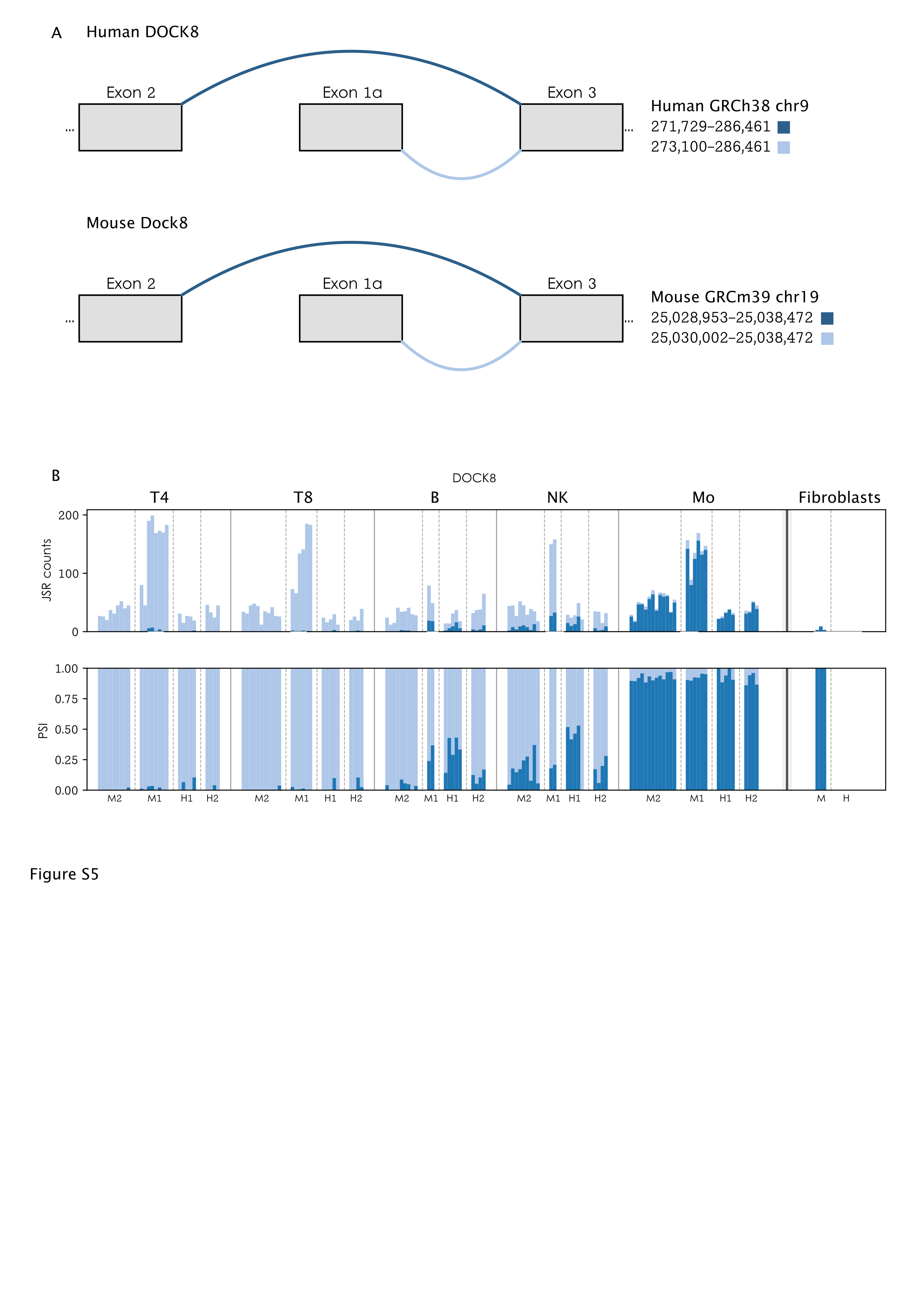

### Image 6.TIFF

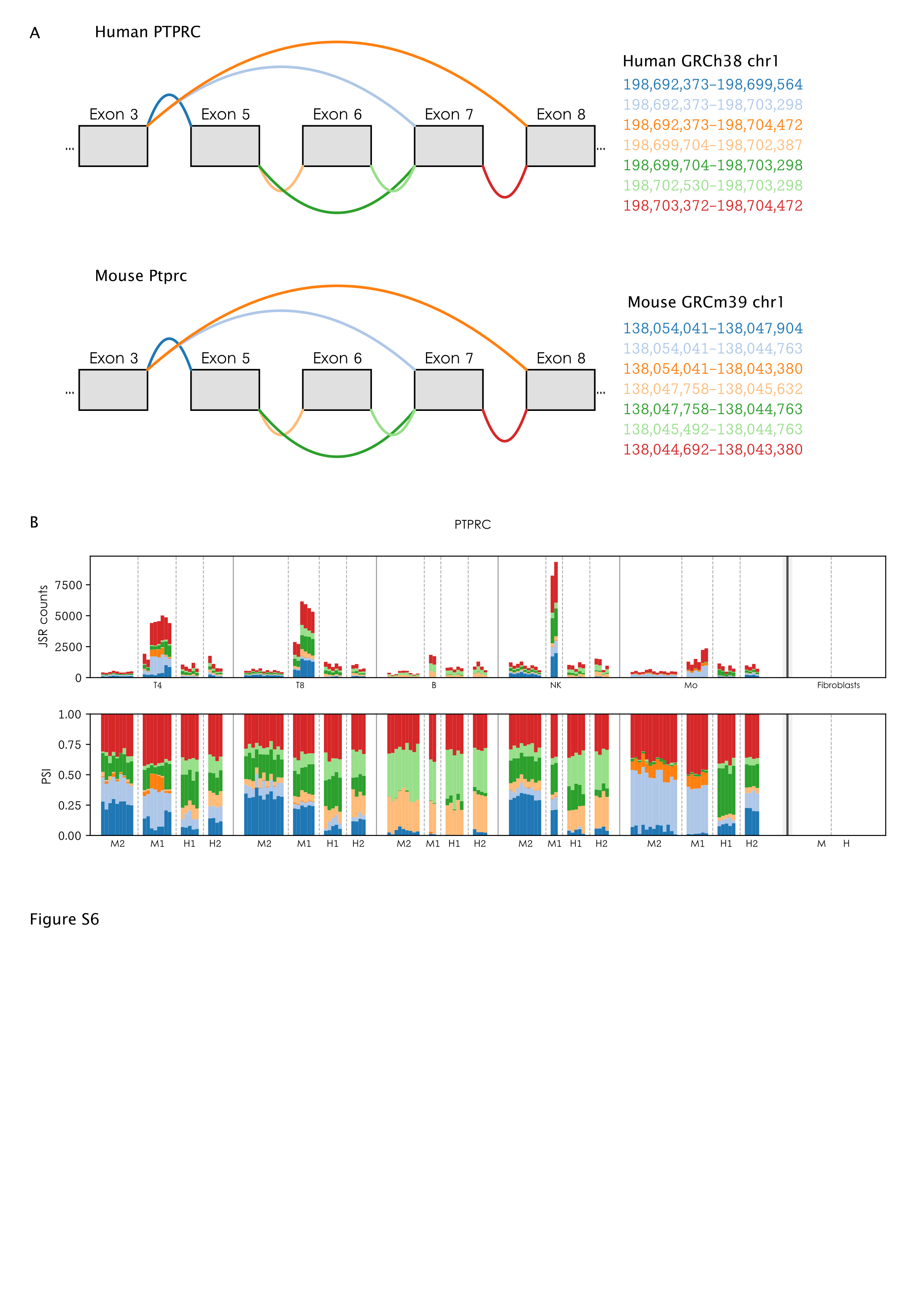

### Image 7.TIFF

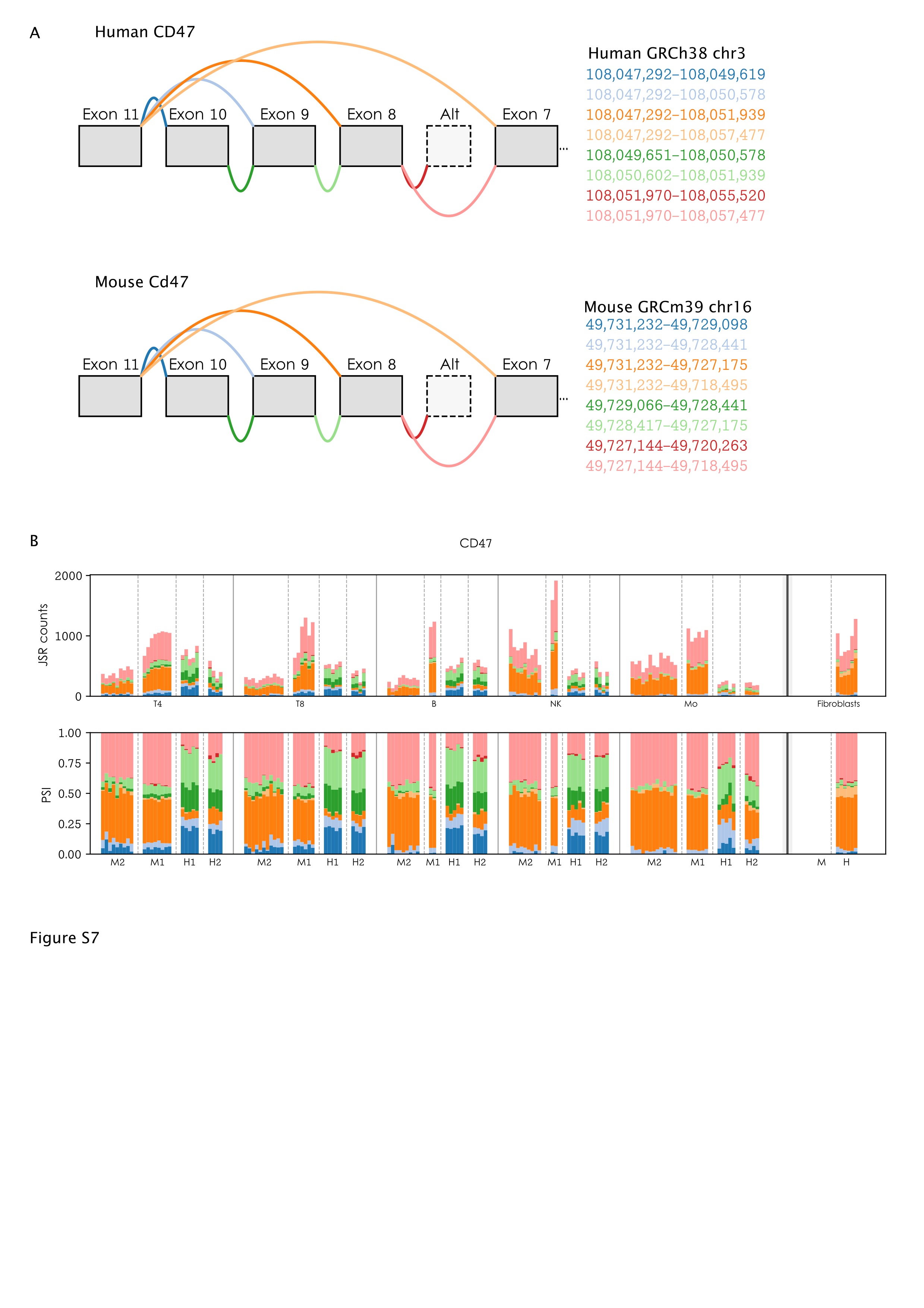

### Image 8.TIFF

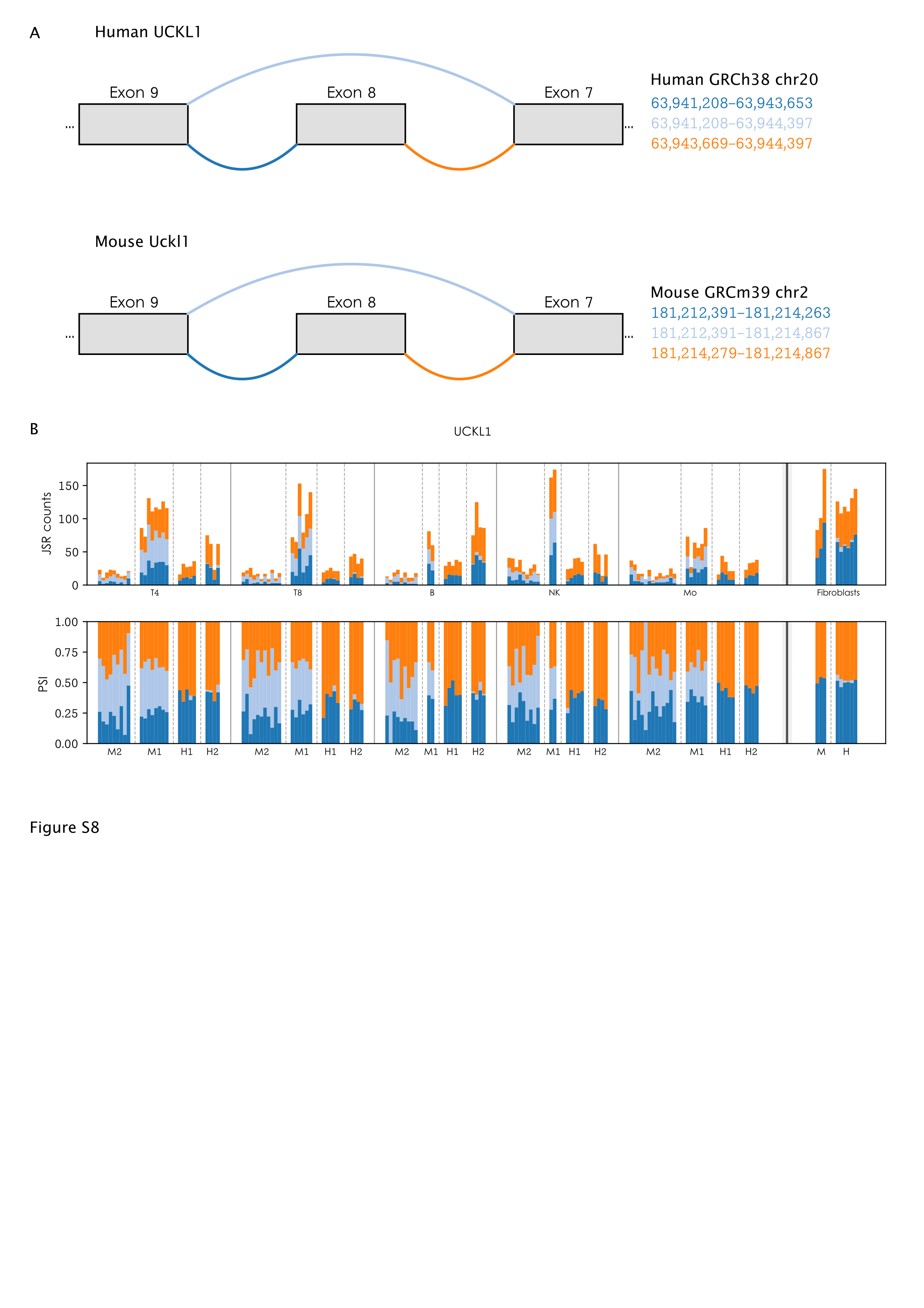
